# Molecular counting of myosin force generators in growing filopodia

**DOI:** 10.1101/2024.05.14.593924

**Authors:** Gillian N. Fitz, Matthew J. Tyska

**Affiliations:** Department of Cell and Developmental Biology, Vanderbilt University School of Medicine, Nashville, TN 27232, U.S.A.

**Keywords:** Myosin, Filopodia, Force

## Abstract

Animal cells build actin-based surface protrusions to enable biological activities ranging from cell motility to mechanosensation to solute uptake. Long-standing models of protrusion growth suggest that actin filament polymerization provides the primary mechanical force for “pushing” the plasma membrane outward at the distal tip. Expanding on these actin-centric models, our recent studies used a chemically inducible system to establish that plasma membrane-bound myosin motors, which are abundant in protrusions and accumulate at the distal tips, can also power robust filopodial growth. How protrusion resident myosins coordinate with actin polymerization to drive elongation remains unclear, in part because the number of force generators and thus, the scale of their mechanical contributions remain undefined. To address this gap, we leveraged the SunTag system to count membrane-bound myosin motors in actively growing filopodia. Using this approach, we found that the number of myosins is log-normally distributed with a mean of 12.0 ± 2.5 motors [GeoMean ± GeoSD] per filopodium. Together with unitary force values and duty ratio estimates derived from biophysical studies for the motor used in these experiments, we calculate that a distal tip population of myosins could generate a time averaged force of ∼tens of pN to elongate filopodia. This range is comparable to the expected force production of actin polymerization in this system, a point that necessitates revision of popular physical models for protrusion growth.

**SIGNIFICANCE STATEMENT:** This study describes the results of in-cell molecular counting experiments to define the number of myosin motors that are mechanically active in growing filopodia. This data should be used to constrain future physical models of the formation of actin-based protrusions.

## INTRODUCTION

Filopodia are fingerlike, micron-scale, membrane-wrapped protrusions that extend from the surface of diverse eukaryotic cell types (1). Structural support for these features is provided by a cytoskeletal core containing of 10-30 actin filaments, cross-linked by the actin bundling protein fascin (2) and uniformly oriented with their fast growing barbed-ends against the plasma membrane at the filopodial tip. Filopodia are found in a range of physiological scenarios, where they enable cells to physically and biochemically interact with neighboring cells and the external environment. In tissues as diverse as mesenchymal cells and neurons, filopodia are critical for cell migration, cell-cell contacts, and wound healing (3-5). Despite their widespread appearance, important questions surrounding the physical basis of filopodial formation remain unanswered.

Cell biological studies and mathematical models dating back several decades suggest that mechanical force generated by the elongation of actin filaments drives the outward deformation of membrane required for filopodial formation (6-8). Here, a balance between the force generated by the fast-growing, barbed ends of actin filaments (F_BE_) and the mechanical barrier provided by tension in the overlying plasma membrane (F_MEM_) dictates whether a distal tip elongates (F_BE_ > F_MEM_), stalls (F_BE_ = F_MEM_), or retracts (F_BE_ < F_MEM_). However, given the actin-rich nature of filopodia and related structures such microvilli and stereocilia, members of the myosin superfamily of actin-based force generators are abundant protrusion residents that could shift the balance of polymerization and membrane forces (9-11). In vertebrates, myosins that contain MyTH4-FERM cargo binding domains offer specific examples of protrusion resident myosins, including Myo10 in filopodia (12), Myo7B in microvilli (13), and Myo15A and Myo7A in stereocilia (14, 15). The functions of these motors are ancient and likely conserved given that myosins with these structural motifs also contribute to filopodial formation in lower eukaryotes (16). Previous studies established that MyTH4-FERM myosins use the core actin bundle as a track to drive tipward transport of cargoes that are needed for protrusion growth and long-term stability (1). Myo10 is a highly studied case; as a processive motor that accumulates at filopodial tips, this motor delivers protein and lipid cargoes that promote filopodia formation, stability, and anchoring (12, 17-22). Indeed, a recent screen using the C-terminal FERM domain of Myo10 as bait identified numerous proteins with wide ranging functions in signaling and regulating cytoskeletal dynamics (17).

In addition to carrying cargo via their C-terminal tails, protrusion resident myosins also interact with the plasma membrane, either directly or indirectly (1), suggesting that these motors could impact the force balance that controls protrusion elongation. Our group recently tested this concept using a genetically encoded, chemically inducible system, which enabled precise temporal control of myosin motor domain recruitment to the plasma membrane. Activation of this system led to rapid and robust elongation of a large number of filopodia (23). This effect was supported by motor domains from structurally distinct MyTH4-FERM domain containing myosins, different membrane binding motifs, and was operational in distinct cell types. These experiments revealed that, beyond powering the tipward transport of cargoes, myosin motors can also contribute to protrusion growth by applying barbed-end-directed force to the overlying plasma membrane, and potentially shifting the balance of force to favor filopodial growth (F_BE_ > F_MEM_). We now refer to this myosin-powered, inducible filopodia system as ‘filoForm’, a name inspired by the plant species *filiformis*, which generates large numbers of long, thin protrusive structures on its leaves; this nomenclature also links to previous filopodia-themed resources adopted by the cytoskeleton field (24-26).

The question of how myosin-generated force promotes protrusion growth remains open. One possibility is that applying tipward force to the plasma membrane increases actin polymerization efficiency by making the fast-growing barbed ends of filaments more accessible for monomer incorporation. Such an effect could be mediated by driving the tipward flow of lipids, which might decrease local membrane tension within the filopodial tip compartment. Alternatively, myosin-generated force might accelerate retrograde flow of the core actin bundle, which could, similarly, make the barbed ends of core bundle actin filaments more accessible, allowing for more efficient polymerization. Although these ideas are intuitive, the mechanistic details of how membrane-bound myosins promote protrusion growth remain unclear due to the lack of quantitative information on the number of force generators (actin vs. myosin) that are active in individual protrusions during growth due to imaging limitations.

To address this gap, we leveraged our ability to initiate the formation of filopodia with filoForm in combination with a single molecule imaging strategy to directly count the number of myosin motors in individual, newly formed filopodia. Using this approach, we found that the number of motors that accumulate in the distal tip compartment during filopodial elongation was highly variable, even across the surface of an individual cell, with a mean of ∼12 motors. Using this value to estimate time-averaged force production, a myosin population of this scale could generate ∼tens of pN, comparable to the expected force production by elongating barbed ends of core bundle actin filaments. The results reported here will be critical for constraining physical models that consider mechanical contributions from both force generating systems: actin filaments and protrusion resident myosins.

## RESULTS

### Combining SunTag_18x_ and filoForm enables counting of individual myosin molecules in newly formed filopodia

To count myosin force generators in elongating filopodia, we first sought an approach that would enable visualization and tracking of single myosin molecules within individual protrusions. To this end, we took advantage of the SUperNova (SunTag) system, a tandem GCN4 repeat scaffold that recruits up to 24 copies of GFP (single-chain variable fragment scFV fused with superfolder GFP) through antibody-peptide interactions (27). The SunTag greatly enhances the fluorescence signal emitted from single tagged molecules, such that conventional imaging methods, including spinning disk confocal microscopy (SDCM), can be used for visualization. Since its creation, the SunTag system has been used to visualize a range of dynamic cellular processes in live cells, including the motion of single kinesin motors along microtubules in U2OS cells, synaptic vesicle transport in cultured neurons, and gene translation in living *Drosophila* embryos (28-30). Our intent in the current study was to combine the high signal/noise ratio offered by the SunTag with the inducible myosin-driven system of filopodial growth, filoForm, described in our previous work (**Fig. 1A**) (23). FiloForm is a genetically encoded system that takes advantage of the rapalog-inducible heterodimerization of FRB and FKBP to dock force-generating myosin motors to the plasma membrane via a membrane-binding motif, CDHR2TM (**Fig. 1B**). In the minutes that follow activation with rapalog, cells undergo robust filopodial elongation. In combining SunTag and filoForm (SunTag-filoForm), our goal was to activate filopodial growth and simultaneously monitor the number of myosin motors that accumulate in growing protrusions (**Fig. 1C**).

**Figure 1:**
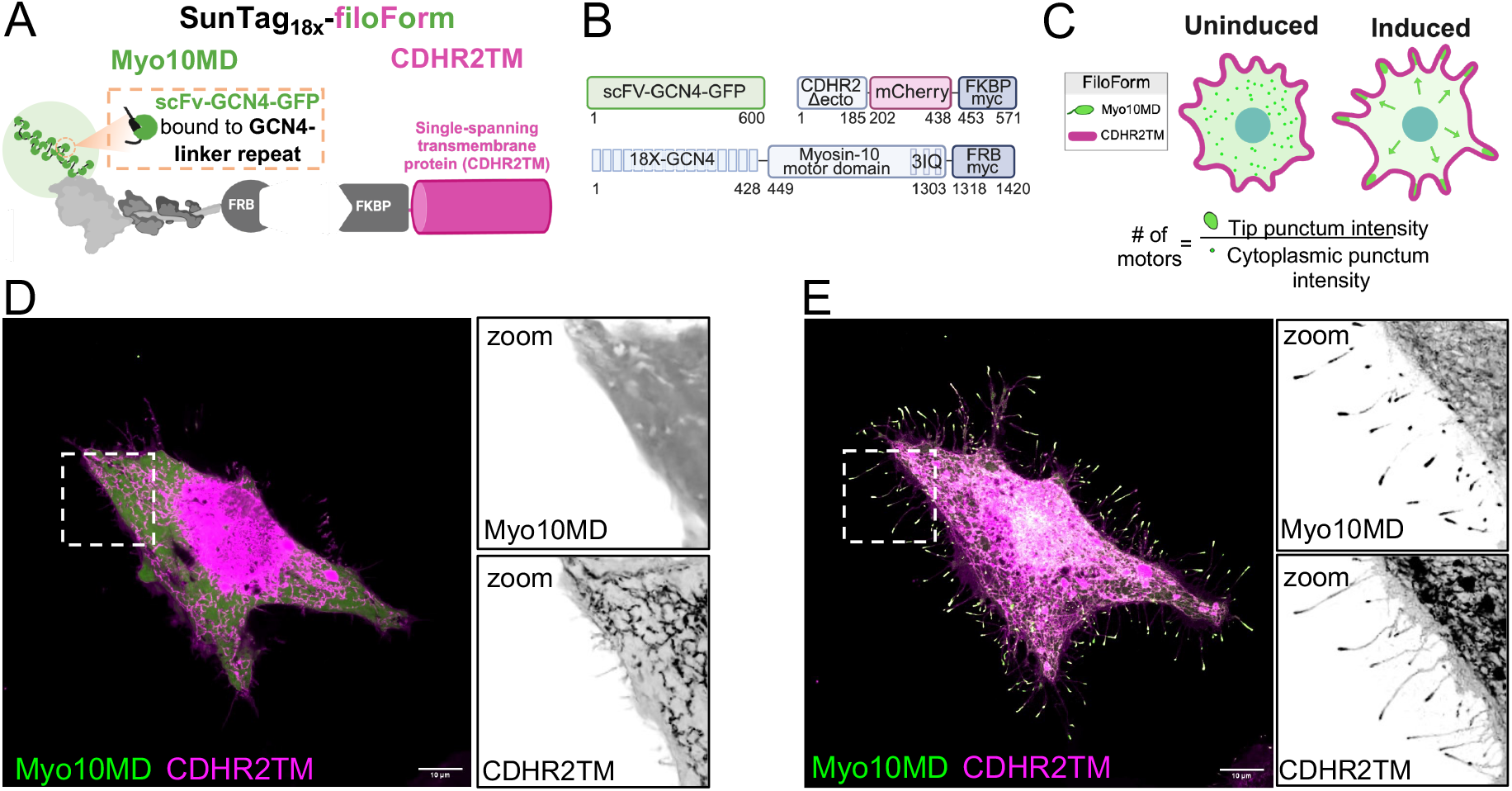
Single molecule confocal imaging using SunTag_18x_-filoForm. (**A**) Schematic of the antibody-peptide labeling strategy (SunTag_18x_) to amplify the signal of single Myo10MD molecules combined with the inducible filoForm system, where FRB-FKBP interactions dock the motor to the membrane anchor (CDHR2TM) upon the addition of rapalog. (**B**) Cartoons depicting the three constructs that comprise SunTag_18x_-filoForm that were transfected into cells to induce filopodia elongation; numbers represent amino acids. (**C**) Graphic of the functional output of these experiments; Myo10MD signal intensity at induced filopodia tips was calibrated by the intensity of cytoplasmic single molecule Myo10MD punctum to calculate the number of motors needed to induce filopodial growth. (**D**) Confocal maximum intensity projection image of a HeLa cell expressing the SunTag_18x_-filoForm system before rapalog addition. (**E**) Confocal maximum intensity projection image of a HeLa cell expressing SunTag_18x_-filoForm 30 min after filopodial induction. Scale bar = 10 μm.

Our previous experiments with filoForm established that myosin-driven filopodial growth is supported by motors with diverse biochemical properties, from classes 3, 10, and 15 (23). For the current studies, however, we focused specifically on myosin-10 motor domain (Myo10MD) as a model myosin force generator. Because the larger variants of the SunTag (i.e. with many tandem GCN4 repeats) are predicted to be bulky and hold some risk of impairing function, we first set out to determine the shortest GCN4 linker that would still enable single molecule visualization using spinning disk confocal microscopy (SDCM), while also supporting filopodial elongation. Testing a range of constructs composed of increasing numbers of tandem GCN4 repeats (4x, 8x, 12x, 14x, 16x, 18x, 20x, and 24x) (**Fig. S1**) revealed that the 18xGCN4 tag (SunTag_18x_) offered the brightest, visible cytoplasmic puncta that still demonstrated robust accumulation in growing filopodia (compare **Fig. 1D** to **Fig. 1E**).

### Defining the fluorescence intensity of a single Myo10MD molecule

HeLa cells transfected with SunTag_18x_-Myo10MD-FRB (referred to herein as Myo10MD) were subject to volume imaging with SDCM, which enabled us to capture filopodia in their entirety. We first sought to measure the intensity of individual Myo10MD molecules in the cytoplasm prior to inducing filopodia elongation with rapalog addition. To measure single Myo10MD molecule intensities unaffected by diffusive motion, we acutely depleted cells of ATP, which is expected to lock myosin motors in a long-lived (i.e. rigor-like) actin-bound state (compare **Fig. 2A** with **Fig. 2B**). To gradually deplete cytoplasmic ATP, we treated Hela cells with 0.05% sodium azide and 10 mM 2-deoxy-D-glucose (31) and then imaged the ventral cell surface using SDCM. Within 15 minutes of ATP depletion, new Myo10MD molecules landed and dwelled in the imaging plane, presumably reflecting strong binding to basal actin stress fibers (compare **Fig. 2C** with **Fig. 2D, Video S2**). Using TrackMate (32), we quantified the sum pixel intensities of single Myo10MD molecules that entered and remained in focus for at least 15 frames (discarding transient events), and then calculated the mean sum intensity per molecule on a per cell basis (28, 32). Quantification of background subtracted Myo10MD intensities revealed that single molecules exhibited a mean of 320 12-bit gray values (**Fig. 2E**). This intensity value was subsequently used as a calibration factor for counting Myo10MD motors in growing filopodia as described below.

**Figure 2:**
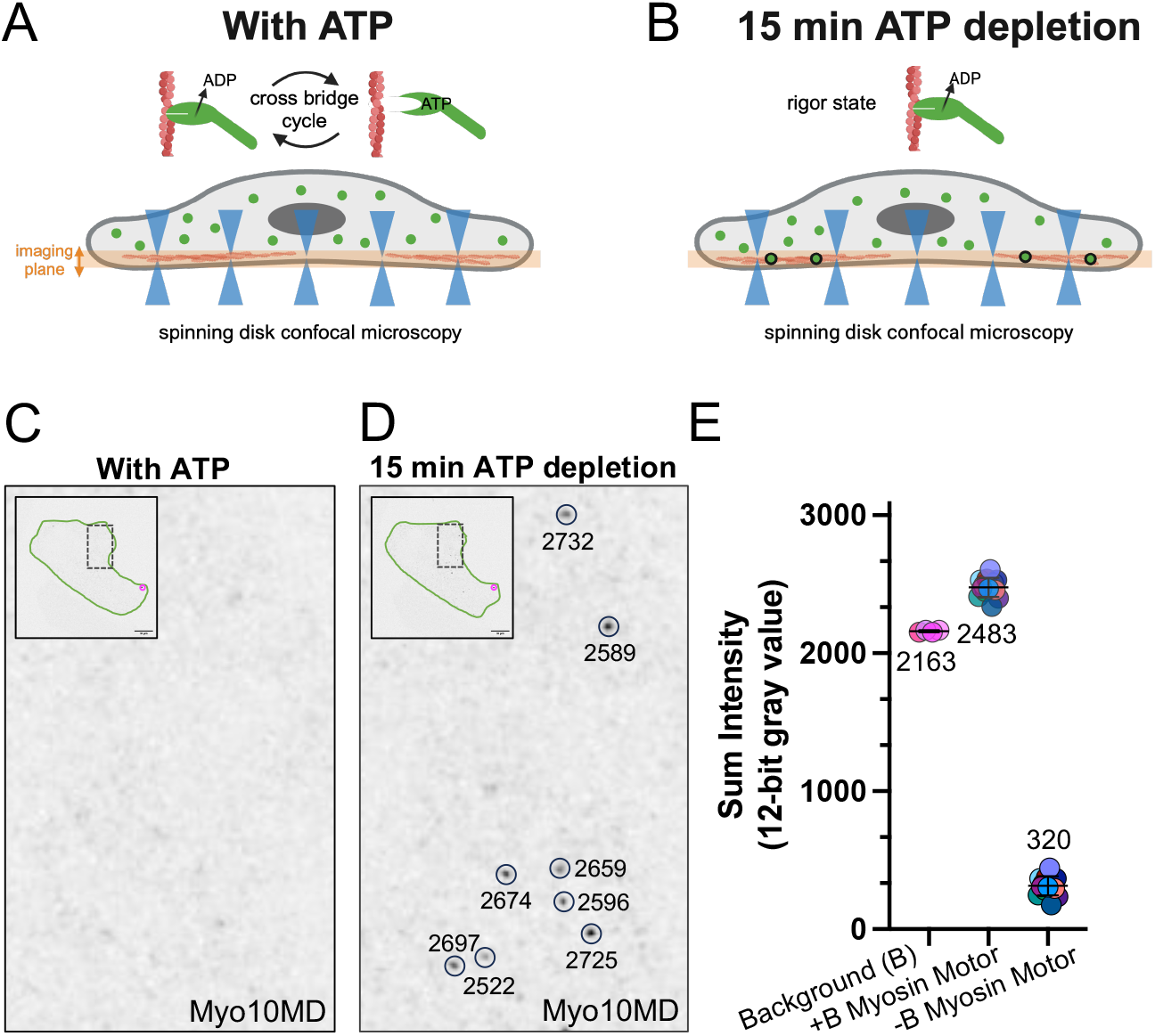
Calibrating the fluorescence intensity of single Myo10MD molecules. (**A and B**) Schematic of ATP depletion experiments used to immobilize Myo10MD molecules. Upon ATP depletion, myosin motors become strongly bound to actin in a rigor state, which increases the number of motors in a single z-plane (A and B, orange box). (**C**) Zoomed image of the ventral surface of a HeLa cell expressing Myo10MD (inverted; black) before ATP depletion; inset shows the cell body outlined in green. (**D**) Zoomed image of the ventral surface of a HeLa cell expressing Myo10MD (inverted; black) 15 min after ATP depletion. Cell body is outlined in green; black circled puncta denote single myosin molecules; eight puncta are circled and shown with the measured sum intensities. (**E**) Sum intensities of Myo10MD molecules from ATP depleted cells; each dot represents the average sum intensity of all puncta measured per cell. n = 100 puncta from 4 cells (background), n = 292 puncta from 13 cells (myosin motor). Values represent the average intensity of background, myosin motor puncta with background included (+B), and myosin motor puncta with background subtracted (-B), with 320 denoting the mean sum intensity of a single tagged myosin motor. Puncta observed pre- and post-ATP depletion (circled in red in C and D) were excluded from the analysis. Scale bar = 10 μm.

### Newly elongated filopodia contain tens of Myo10MD molecules

To quantify the number of Myo10MD motors that accumulate in newly formed filopodia, we induced filopodia elongation using SunTag_18x_-filoForm, performed volume imaging with SDCM, and used 3D thresholding to quantify the sum intensity of motor signal at the distal tips, 30 min after rapalog activation. Using the single molecule calibration value determined above, we found that filopodia contained ∼12 myosin motors per protrusion (12.0 ± 2.5 motors, GeoMean ± GeoSD), although induced protrusions demonstrated significant variability, containing as few as 2 and as many as 349 motors (**Fig. 3A**). We were curious to determine how these values compared to protrusions generated by the overexpression of wild-type Myo10, which is known to drive robust filopodial initiation and elongation (12). To test this, we created a full-length Myo10 (Myo10FL) construct tagged with SunTag_18x_ (**Fig. 1**). We transfected HeLa cells with SunTag_18x_-Myo10FL, and again used SDCM to examine motor accumulation at filopodial tips. Given the nature of conventional transient transfection experiments, the population of protrusions that we analyzed in this case contained both nascent and mature filopodia. 3D thresholding and intensity analysis revealed that filopodia tips in Myo10FL expressing cells contained ∼9 myosin motors per protrusion (**Fig. 3B**), comparable to what we observed using the filoForm described above. Thus, independent of the mode of myosin-dependent filopodia assembly (induced vs. stochastic), filopodia contain ∼tens of motors (**Fig. 3B**).

**Figure 3.**
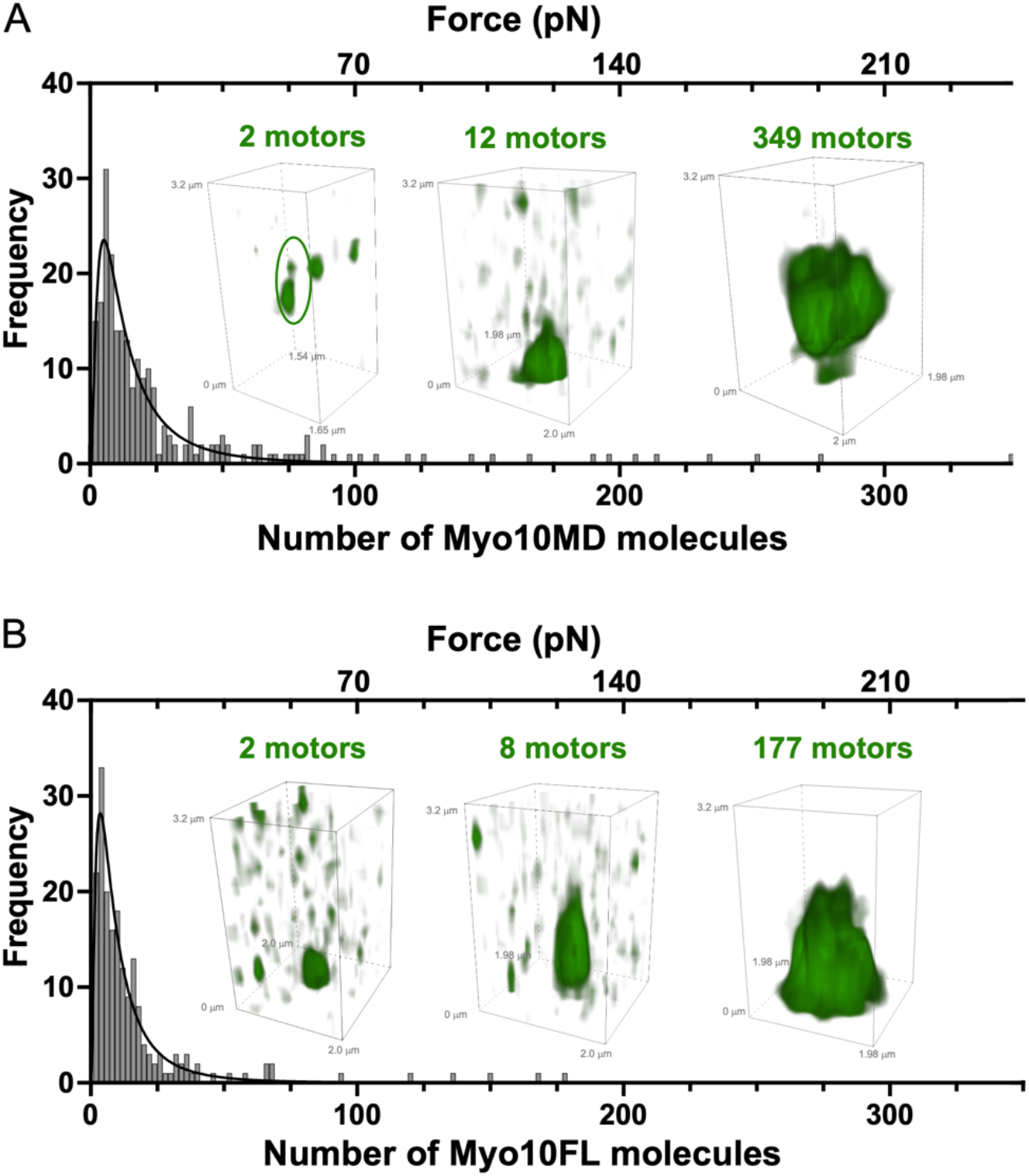
Newly formed filopodia contain tens of Myo10MD molecules. (**A**) Distribution of the number of Myo10MD molecules counted at the tips of newly formed filopodia using SunTag_18x_-filoForm; curve fit represents a log-normal distribution with 12.0 ± 2.5 motors (GeoMean ± GeoSD). Green inset images depict deconvolved volume projections of filopodial tips showing a range of motor numbers; n = 14 cells, 237 individual filopodial tips. (**B**) Log-normal distribution of the number of Myo10FL molecules at filopodial tips 24 hrs post transfection; curve fit represents a log-normal distribution with 8.6 ± 2.60 motors (GeoMean ± GeoSD). Green inset images depict deconvolved volume projections of filopodial tips showing a range of motor numbers; n = 18 cells, 189 individual filopodia. For both **A** and **B**, top x-axis represents the correlating amount of force calculated using a duty ratio of 0.7.

### Filopodia dynamics are controlled by the number of tip localized Myo10MD molecules

Next, we wondered if the dynamic properties of growing filopodia are impacted by the number of myosin motors that accumulate at the distal tips. To investigate this possibility, we imaged the elongation of nascent filopodia at low z-resolution for 30 min, and then acquired high-resolution volumes to enable the counting of myosin motors as described above (**Fig. 4A**). We then generated trajectories from distal tip motion during elongation to calculate the elongation rate, persistence of elongation, and maximum final length for each filopodium. Trajectories were then binned into low and high motor groups depending on whether the number of motors at the distal tip was below or above the GeoMean of 12 (from **Fig. 3A**). Filopodia in the high motor group elongated faster, were more persistent, and reached longer final filopodia lengths (**Fig. 4B, 4C, and 4D**). While we noted a significant increase in both tip velocity and persistence, the most significant increase was observed in the final length of filopodia. These results suggests that force generated by membrane-bound myosins delays the stalling of growth that ultimately limits filopodia length.

**Figure 4.**
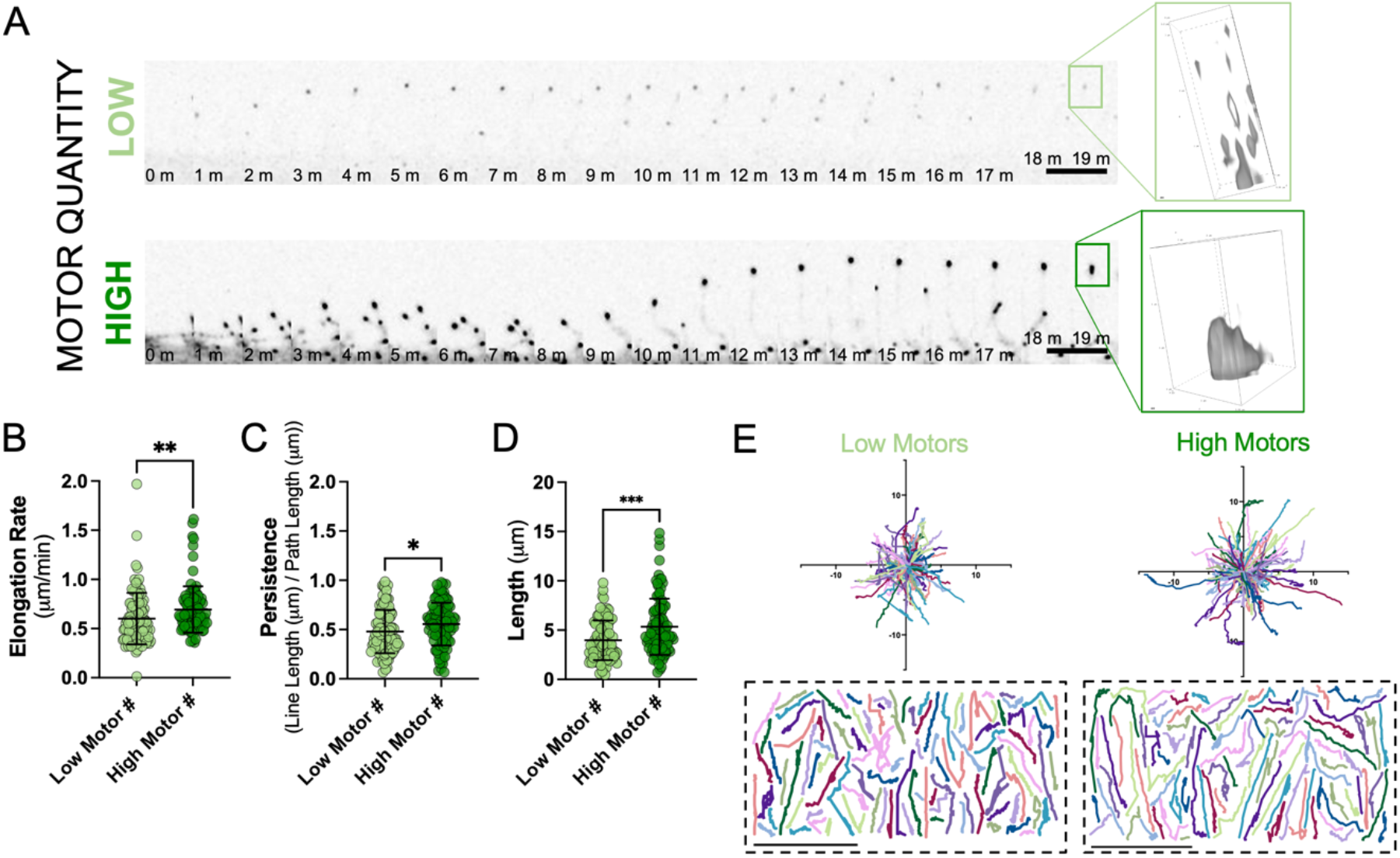
Filopodial growth dynamics depend on the number of tip targeted myosin motors. (**A**) Montage of a individual filopodia elongating from a HeLa cell expressing SunTag_18x_-filoForm containing a low (2; top) or high (47; bottom) number of motors. Right image depicts a deconvolved volume projection of the elongating filopodial tip at the final time point. Scale bar = 5 μm; LUTs matched to high motor montage. Quantification of (**B**) elongation rate, (**C**) persistence of elongation, and (**D**) final length for filopodia containing low (< 12; light green) or high (> 12; dark green) numbers of motors. Error bars represent the mean ± SD; * p ≤ 0.05, ** p ≤ 0.05, *** p ≤ 0.001. (**E)** Rose plots of individual trajectories of filopodial tips containing low (< 12; light green) or high (> 12; dark green) numbers of motors. Dashed boxes below depict the individual filopodial tracks; n = 100 (low), 102 (high) filopodia (B-D). Scale axis = 10 μm.

## DISCUSSION

Our recent work established that membrane-bound myosins can tilt the force balance at filopodial tips to promote elongation, presumably by potentiating the force generated by core bundle actin filaments (23). To define the scale of myosin motor force production, we combined the SunTag with the filoForm system, which allows for both single molecule visualization and inducible control of filopodia formation via the docking of myosin motors to the plasma membrane. The primary advance offered by our approach is that measurements are time-resolved, allowing us to focus our analysis on actively growing filopodia immediately after induction of the filoForm system.

Monitoring myosin accumulation in newly grown filopodia revealed that only ∼12 myosin motors are needed to induce the rapid elongation of protrusions. When we overexpressed a SunTagged variant of Myo10FL and examined intensities at filopodia tips in live cells ∼24 hours post transfection, we obtained a similar result with a mean of ∼9 motors per filopodium. Although we know with certainty the age of nascent protrusions that grow after activating the filoForm system, we are unable to know the age of filopodia formed by full length Myo10 given the stochastic nature of protrusion growth in those experiments. Nevertheless, filopodia in the latter case are expected to be older on average relative to induced protrusions. Thus, filopodial aging (or maturation) does not appear to significantly change the number of myosin motors in the distal tip compartment. This point further implies that the number of myosins promoting elongation may be set at the initiation of filopodial growth. In contrast to our findings, a recent study in U2OS cells demonstrated that over-expression of Myo10FL leads to filopodial tips that contain hundreds of motors (median of ∼360 molecules), with a large range around this value (2-65,000 molecules) (33). Although we employed a direct single molecule visualization strategy, that study used a population level biochemical approach combined with epifluorescence microscopy to approximate the number of motors in individual cells, and in turn, single filopodia. Interestingly, the authors did describe a population of filopodia characterized by dim tip puncta, representing only tens of Myo10 motors, which would be generally consistent with the results we present here.

How much mechanical force could ∼tens of myosin motors generate at the tip of a filopodium? The time-averaged force (F_avg_) produced by a population of motors is related to the number of force generators in the population (in this case ∼12), the unitary force produced by individual motors (F_uni_ ∼1 pN), and the duty ratio, i.e. the fraction of the total ATPase cycle spent in a force-generating state (F_AVG_ = F_UNI_ × # of motors × duty ratio) (34). Duty ratio estimates for the Myo10 motor used in our studies are wide ranging (0.16-0.7) (35, 36), but the higher value of 0.7 yields F_AVG_ of ∼8.4 pN (35). Notably, this calculation does not consider the lifetime of the membrane-bound state for force generating myosins, which is expected to be long for motors activated in the filoForm system, given the strong binding interactions between the rapalog and FKBP/FRB modules (37). Because native protrusion myosins probably bind/unbind the membrane at a higher frequency, our calculation of F_AVG_ derived above may represent an overestimate.

How does F_AVG_ generated by tip targeted myosins compare to the force generated by elongating actin filaments in the filopodium (F_BE_)? Previous biophysical studies established that polymerizing actin filaments exert up to ∼1 pN on an opposing surface (38, 39). Filopodia core bundles assembled in neurons contain up to ∼20 filaments (40), putting an average maximum force generated by that mechanism at ∼20 pN. However, growing barbed-ends of a parallel bundle are unlikely to exhibit synchronous and continuous insertion of new actin monomers (41), and therefore the number of filaments exerting force on the plasma membrane at any point in time will likely be a fraction of the total number. With this in mind, the level of force generated by actin and membrane-bound myosins during filopodial elongation may be comparable, i.e. within the same order of magnitude.

The number of myosin motors that accumulate at the tips of newly grown filopodia was highly variable across the surface of a single cell (**Fig. S2**). What does this variability suggest about factors that control protrusion growth? Based on popular models of filopodia formation, in-plane tension in the plasma membrane is viewed as the primary physical barrier that growing actin filaments must contend with to elongate these structures (42, 43). In the context of our assays, membrane tension could vary from cell to cell (e.g. based on overall morphology or surface attached area) or across the surface of an individual cell. Consistent with this latter possibility, previous biophysical studies established that membrane tension does vary locally, with single cells yielding a wide range of tether forces (∼5-30 pN) depending on where the tether was pulled (43-45). From this perspective, the large variability in distal tip myosin accumulation within individual cells might reflect variability in local membrane mechanics, with regions of high and low tension across the surface requiring higher or lower levels of myosin force production, respectively, for filopodial elongation. Another possibility relates to inherent variation in the local accumulation of other factors that promote filopodia growth. This idea is supported by a recent study from Gallop et al., which revealed striking variability in the composition of filopodial tip complexes and further highlighted that stochastic accumulation of different combinations of factors are able to support filopodial growth (46).

While generating and optimizing SunTagged myosin motor constructs, we noted that the 24xGCN4 tag (SunTag_24x_) was unable to elongate filopodia. One trivial explanation is that this long linker variant was unable to acquire a stable fold, and therefore remained non-functional in the cytoplasm. However, a more intriguing possibility is that the long linker variant was too bulky to enter the tight confines of the space between the actin core and the encapsulating plasma membrane, at the filopodial base. Although a size exclusion limit has not yet been reported for filopodia, we examined solved structures and AlphaFold predictions to estimate the volumes occupied by the tagged myosin constructs used in our studies (**Fig. S3**) (47, 48). The volumes of a single Myo10MD tagged with FRB, a single EGFP, and a single scFV bound to a GCN4 linker, are ∼130 nm^3^, ∼29 nm^3^, and ∼72 nm^3^, respectively. Extrapolating these values leads to approximate volumes for EGFP-Myo10MD-FRB, 20xGCN4-Myo10MD-FRB and 24xGCN4-Myo10MD-FRB of ∼159 nm^3^, ∼1568 nm^3^, and ∼1855 nm^3^, respectively. Are these molecular volumes too large to enter the space between the actin core bundle and overlying plasma membrane at the base of a newly forming filopodia? Cryo-TEM images of *Dictyostelium* filopodia offer an estimate of the distance between the core actin bundle and encapsulating membrane of ∼15 nm (49), and a sphere with this diameter would occupy ∼1767 nm^3^. Using this value as a molecular volume threshold, the 20xGCN4-Myo10MD-FRB construct (∼1568 nm^3^), but not the 24XGCN4-Myo10MD-FRB construct (∼1855 nm^3^), would be expected to accumulate in filopodia, which matches our experimental results. These results suggest that access to a filopodium may be restricted by the size of the opening at its base. Future studies working with native filopodial resident proteins will need to explore this possibility further.

In conclusion, our investigations revealed that newly forming protrusions only require tens of myosin motors to elongate. A population of this scale could generate force (F_AVG_) at a level comparable to that produced by polymerizing filament barbed ends (F_BE_), and therefore make a significant contribution toward overcoming the physical barrier that impedes filopodial growth (F_MEM_). The large variability in motor numbers between protrusions further reinforces the previously described mosaicism of mechanisms that drive protrusion growth (46). Importantly, these findings necessitate updates to popular physical models of filopodial growth, which generally consider polymerizing actin filaments the sole force generators in these structures.

## MATERIALS AND METHODS

### Lead Contact

Further information and requests for resources and reagents should be directed to and will be fulfilled by the lead contact, Matthew J. Tyska.

### Materials availability

Plasmids generated in this study will be made available from the <u>lead contact</u> on request.

### Experimental model and subject details Cell culture

HeLa cells were cultured at 37ºC and 5% CO_2_ in Dulbecco’s modified Eagle’s medium (DMEM) (Corning #10-013-CV) with high glucose and 2 mM L-Glutamine supplemented with 10% fetal bovine serum (FBS).

### Cloning and constructs

All PCR amplification was completed using Q5 High-Fidelity DNA Polymerase (NEB #M0491S) for all PCR amplifications including site-directed mutagenesis.

*Rapalog Inducible (FRB-FKBP) filopodial elongation system*: EGFP-Myo10MD-FRB and the single-spanning transmembrane domain membrane docking construct (CDHRTM-mCherry-FKBP) was generated as previously described (23).

*Myosin-10 Motor Domain GCN4 Linkers*: To generate the GCN4 linkers (4x-, 8x-, 12x-, 16x-, 18x-, 20x-, 24x-) tagged to Myo10MD-FRB, we replaced the EGFP from the previously generated EGFP-Myo10MD-FRB (above) with GCN4 repeats from pHRdSV40-K560-24xGCN4_v4 (addgene #72229) via Gibson assembly (NEB #E2621S) using the following primer sets: (24x) 5’-acccaagctgggccaccatggaagaacttttgagcaagaattatcatcttgagaacgaagtg-3’ and 5’-

attcgagatctgagtccggactttttaagtcgggctacttcattctcgaga-3’, (20x) 5’-

acccaagctgggccaccatggaagaacttttg-3’ and 5’-

-attcgagatctgagtccggacttctttagtcgagccacctcgttctcg -3’, (18x) 5’-

acccaagctgggccaccatggaagaacttttgagcaagaattatcatcttgagaacgaagtg-3’ and 5’-

attcgagatctgagtccggatttcttcaagcgggcgact -3’, (16x) 5’-

acccaagctgggccaccatggaagaacttttgagcaagaattatcatcttgagaacgaagtg -3’ and 5’-

attcgagatctgagtccggattttttgagcctggcgacttca -3’, (14x) 5’-

acccaagctgggccaccatggaagaacttttgagcaagaattatcatcttgagaacgaagtg -3’ and 5’-

attcgagatctgagtccggatttcttcaatctcgcgacctcattc -3’, (12x) 5’-

acccaagctgggccaccatggaagaacttttgagcaagaattatcatcttgagaacgaagtg -3’ and 5’-

attcgagatctgagtccggatttctttaagcgcgcgacttcgttctctaaatgatag -3’, (8x) 5’-

acccaagctgggccaccatggaagaacttttgagcaagaattatcatcttgagaacgaagtg-3’ and 5’-

attcgagatctgagtccggatttctttaatcgagctacttcgttttcgagg -3’, (4x) 5’-

acccaagctgggccaccatggaagaacttttgagcaagaattatcatcttgagaacgaagtg -3’ and 5’-

attcgagatctgagtccggacttttttagccgagccacttcgttt -3’.

*scFV-GFP:* We generated scFV-GFP by removing the nuclear localization sequence via an early stop codon from pHR-scFv-GCN4-sfGFP-GB1-NLS-dWPRE (Addgene #60906) with the following primers: 5’-tggtagctgaggtggtactagtcccaagaagaagc-3’ and 5’-ccacctcagctaccaccaccttcggttacc-3’. The reaction was then digested with DpnI (NEB #R0176S).

### Transfections

Transfections were performed using Lipofectamine 2000 (Thermo Fischer #11668019) according to the manufacturer’s protocol. 2.5 x 10^5^ HeLa cells were resuspended and incubated in Lipofectamine for 16 hrs in 35mm glass-bottom dishes (Cellvis #D35-20-1.5-N) before being rinsed once in 1X PBS and replaced with media and live imaged.

### Drug treatments

To oligomerize FRB and FKBP constructs, transfected cells were treated with 500 nM of A/C Heterodimerizer (Takara #635057) at the onset of live imagining. To deplete ATP for the ATP depletion experiments, transfected HeLa cells were switched to glucose free DMEM (Fischer #11-966-025) supplemented with 2 mM L-glutamine and 10% FBS. Before the onset of live cell imaging, 35 mm dishes were spiked with 0.05% sodium azide (Fischer #11-966-025) and 10 mM 2-deoxy-d-glucose (Sigma #D3179). For live-cell imaging of individual filopodia, 0.5% methylcellulose diluted in DMEM with high glucose and 2 mM L-Glutamine supplemented with 10% FBS was added prior to imaging.

### Light microscopy and image processing

Live-cell imaging was performed on a Nikon Ti2 inverted light microscope equipped with a Yokogawa CSU-W1 spinning disk head, equipped with 405 nm, 488 nm, 561 nm, and 647 nm excitation LASERs, a 100X Apo TIRF 100x/1.45 NA objective, and a Photometrics Prime 95B sCMOS camera. Cells were maintained in a stage top incubator at 37ºC with 5% CO2 (Tokai Hit). For imaging in all microscope modalities, imaging acquisition parameters were matched between samples during image acquisition. For imaging of individual puncta, a single z-stack acquisition was taken for 5 sec at 0.05-0.06 ms/frame prior to depleting ATP. ATP depletion media was then added and 15 min later, the same cells were imaged for 5 s at 0.05-0.05 ms/frame. For imaging the tips of filopodia that had elongated, a 0.5 μm z-stack was acquired, rapalog was added to induce filopodia elongation, and after 30 m the same cell was imaged using a 3.0 μm z-stack to acquire 3D volume tip information. For imaging to quantify the dynamic properties of filopodial elongation based on motor number, transfected cells were imaged every 15 sec for 30 m using a 0.5 μm z-stack, and at the end of 30 m a single, 3.0 μm z-stack was taken. All images were denoised and deconvolved in Nikon Elements. As filopodia are thin structures, LUTs were optimized to facilitate visualization in figures.

### Quantification and statistical analysis

All images were processed and analyzed using Nikon Elements software or FIJI software (https://fiji.sc/).

#### Analysis of individual Myo10MD puncta after ATP depletion

Fiji’s TrackMate was used to quantify the sum intensity of individual puncta. TrackMate was run under the following parameters: all 82 frames were included, DoG detector, 0.5 μm diameter with a 0.75 quality threshold, no initial thresholding restrictions, LAP tracker, 1.0 μm max distance frame linking, allow gap closing with a max distance of 0.5 μm with a max gap frame of 2. Puncta were removed if seen in the intial image prior to ATP depletion, and puncta were only included if they remained in frame for 15 or more frames. An average sum intensity was then taken of the sum intensity of each punctum over the 15 or more frames it appeared. For background puncta, the same sized ROI from TrackMate was used to measure 100 points background sum intensity from each of the 4 separate imaging days. The intensity of a single Myo10MD punctum was then measured as the average of all Myo10MD puncta measured minus the average background intensity.

#### Analysis of the number of motors at the tips of filopodia

Prior to inducing filopodial elongation, we generated a single image with low z-resolution (0.5 μm) of transfected HeLa cells. Transfected HeLa cells were then induced to elongate filopodia, and after 30 min a single image with high z-resolution (3 μm) was captured. A 2 x 2 μm crop of filopodial tips was then taken from any filopodial that had emerged compared to the initial low z-resolution image. This cropped tip was then threshold in using the green, Myo10MD, channel in Nikon Elements to capture the sum intensity of the tip. Using the background pixel intensity from our single puncta analysis, we then quantified the number of voxels (3D pixel) were captured in the 3D tip and subtracted the number of voxels multiplied by the background intensity from the 3D tip. This value was then divided by the background subtracted individual puncta to identify the number of Myo10MDs in each individual filopodial tip. Radial plot of the SunTag_18x_-filoForm was generated using OriginPro.

#### Measuring the dynamics of induced filopodia

Lengths, velocity, and persistence of elongating filopodia were analyzed using “tracking” in Nikon Elements using the Myo10MD tip intensity. Velocity of individual filopodia was determined by dividing the maximum line length by the duration of time it took to reach the maximum length, maximum lengths were measured using line length (length of a straight line from the track origin to the current point), radial tracks were normalized to set the origin to (0,0), and persistence was measured as the line length divided by the path length (sum of the segment distances from the first frame to the current frame) of each individual track. Each individual track was analyzed to include the start of elongation and stopped when a maximum length was reached.

#### Statistical analysis

For the purpose of generating SuperPlots (50) we considered the number of individual cells equivalent to the number of biological replicates; this approach allowed us to build plots that communicate the variability between cells *and* between individual punctum. All experiments were completed at least in triplicate and the number of biological replicates (n) in each case is defined in each figure legend. Statistical significance was performed using the unpaired Student’s t-test for comparisons and the paired Student’s t-test for comparisons of the sum intensities of punctum without and with ATP depletion and for the number of motors at filopodial tips. All statistical analysis were computed using PRISM v.10.0.2 (GraphPad).

## Supporting information

Fitz et al Supp VideoS1 and S2 legend

Fitz et al Supp Figure 3

Fitz et al Supp Figure 2

Fitz et al Supp Figure 1

Fitz et al Video S1

Fitz et al Video S2

## Acknowledgments

The authors would like to thank all members of the M.J.T. laboratory for their feedback and guidance. This work was supported by the NIH NIDDK National Research Service Award F31-DK130599 (G.N.F.) an NIH grants R01-DK125546, R01-DK095811, and R01-DK111949 (M.J.T.).

We also acknowledge the Lacks family and are grateful for the use of HeLa cells, which heavily contributed to the discoveries in this work.

