## Supplementary material for "Molecular counting of myosin force generators in growing filopodia": Fitz et al Supp VideoS1 and S2 legend

**Video S1. Membrane bound 18xGCN4-Myo10MD drives robust elongation of filopodia, related to Figure 1.** Live cell movie of maximum intensity projections of HeLa cell expressing 18xGCN4-Myo10MD-FRB with scFV-GFP (green) and CDHR2TM-mCherry-FKBP (magenta) treated with rapalog. 0.6 μm maximum intensity projections are composed of 4 x 0.2 μm confocal slices.

**VideoS2. ATP depletion increase the number of motor molecules in the imaging plane, related to Figure 2.** Live cell movie of a single HeLa cell expressing 18XGCN4-Myo10MD and scFV-GFP (inverted; black) prior to ATP depletion (left) and 15 minutes after ATP depletion (right). Movie was acquired using a single z-plane focused on the basal surface of the cell. Scale bar = 10 μm. Movies were acquired at no delay (60 ms/frame).
