## Supplementary material for "Molecular counting of myosin force generators in growing filopodia": Fitz et al Supp Figure 3

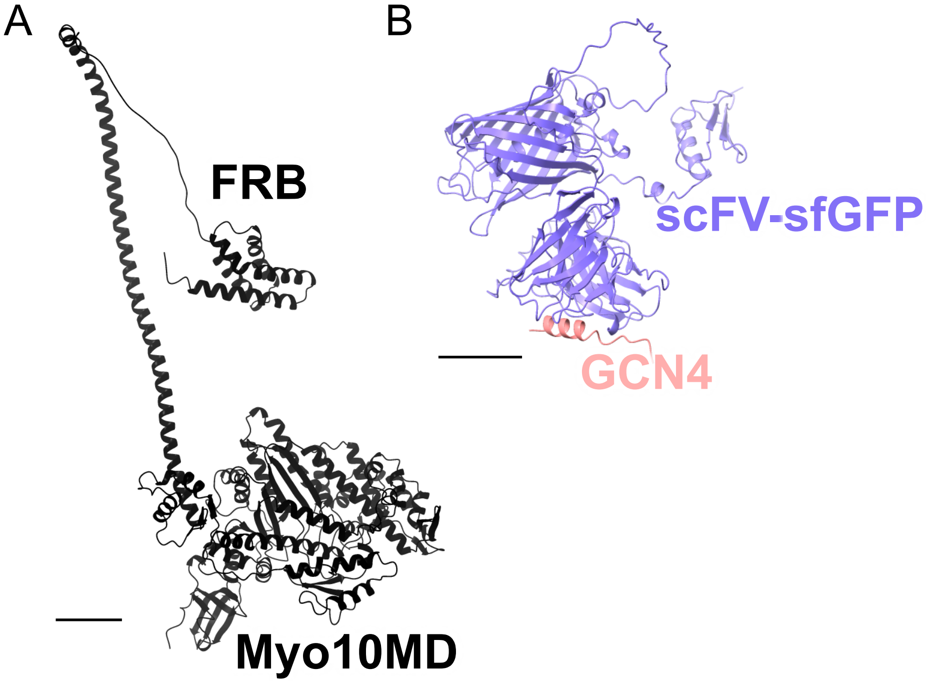


**Fig. S3. AlphaFold predictions used for molecular volume calculations.** (**A**) AlphaFold prediction of the Myo10MD-FRB construct used in these studies without a tag (130 $nm^3$). (**B**) AlphaFold prediction of a single chain variable fragment superfolder-GFP (scFV-sfGFP) (purple) bound to a single GCN4-linker (pink) (72 $nm^3$). Scale bars = 20 Å.
