## Supplementary material for "Molecular counting of myosin force generators in growing filopodia": Fitz et al Supp Figure 2

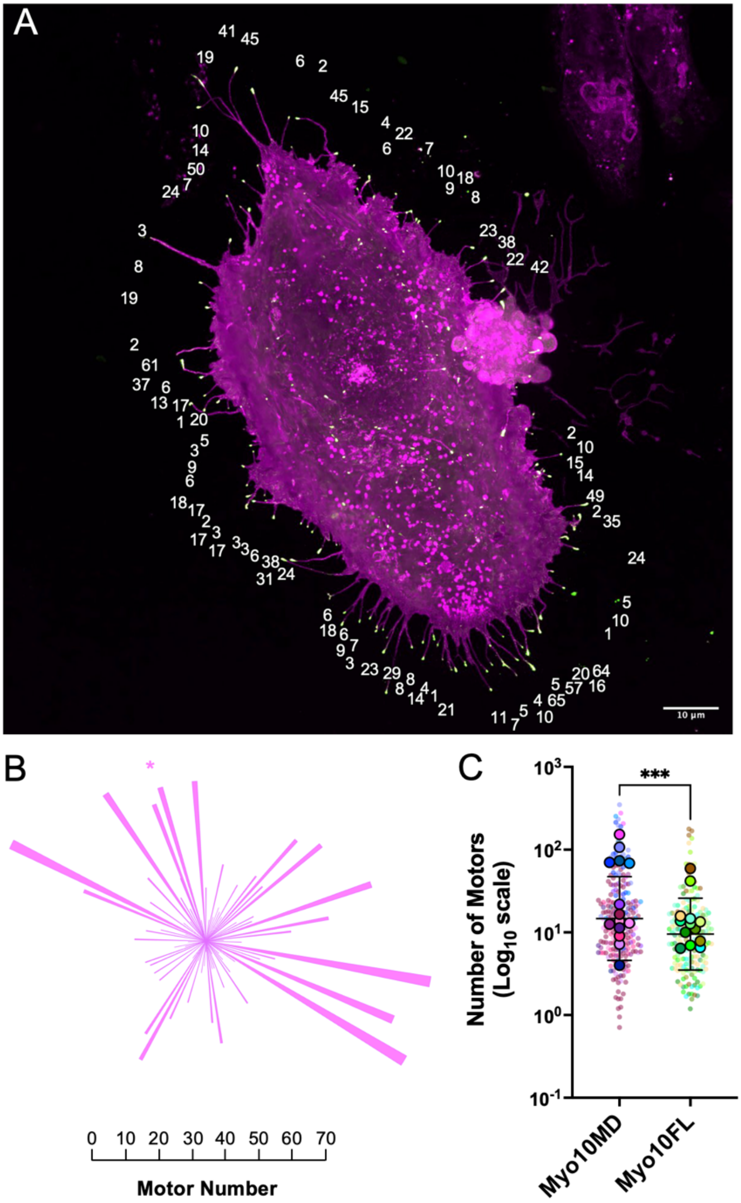


**Fig. S2. Spatial mapping of the distribution of motor numbers at filopodial tips. (A)** Confocal maximum intensity projection image of a single HeLa cell expressing SunTag$_{18x}$-filoForm 30 min after filopodial induction. Values denote the number of Myo10MD molecules at individual filopodial tips. Scale bar = 10 $\mu$m. **(B)** Radial bar plot of the spatial distribution of the number of motors from filopodia shown in A (asterisk denotes the topmost filopodium with 41 motors in A). **(C)** Superplot of the number of myosin motors at the tips of individual filopodia generated with SunTag$_{18x}$-filoForm (Myo10MD) or EGFP-Myosin10-FL (Myo10FL). Error bars represent the mean ± SD; *** $p \leq 0.001$. n = 14 cells, 182 individual filopodial tips (Myo10MD), n = 18 cells, 189 individual filopodia (Myo10FL).
