## Supplementary material for "Molecular counting of myosin force generators in growing filopodia": Fitz et al Supp Figure 1

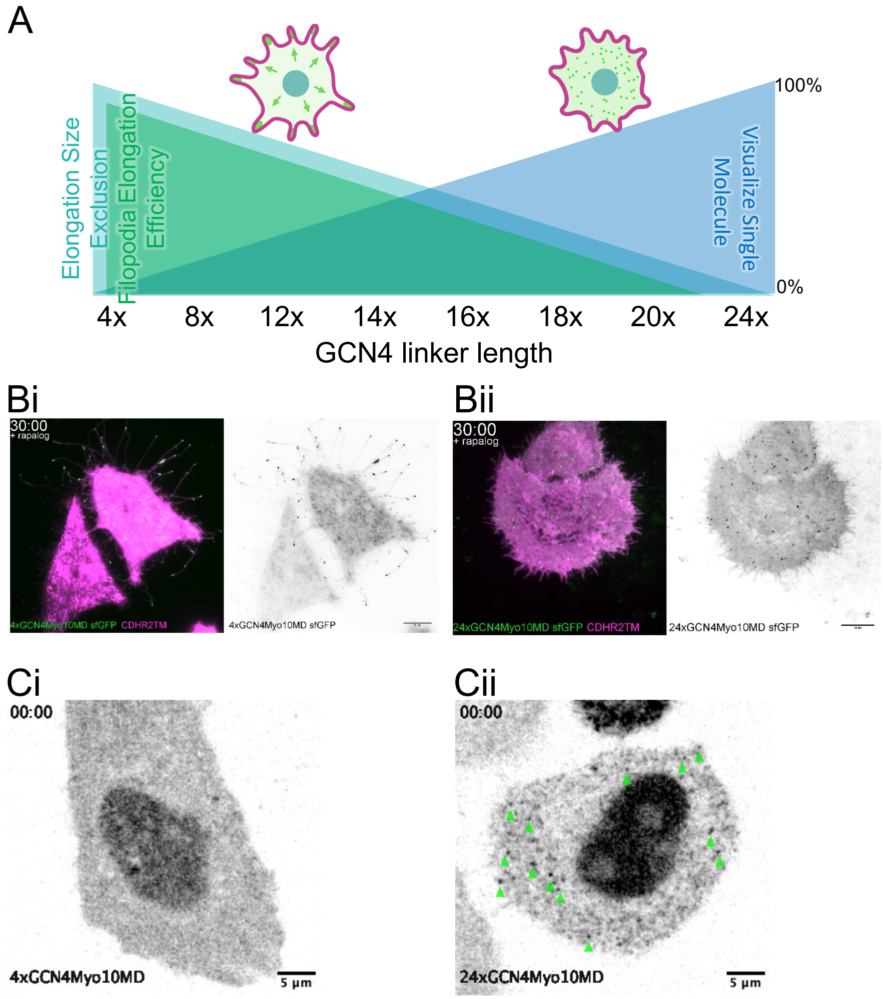


**Figure S1. Optimizing the SunTag-filoForm system.** (**A**) Cartoon schematic of the observed inverse relationship between GCN4 linker length and ability to elongate filopodia following activation with rapalog. (**B**) Spinning-disc confocal microscopy images of single HeLa cells transfected with CDHR2TM (magenta) and either a 4xGCN4-Myo10MD (**Bi**) or 24xGCN4-Myo10MD (**Bii**) linker constructs (green) 30 min after the addition of rapalog; scale bar = 10 μm. (**C**) TIRF images of single HeLa cells transfected with CDHR2TM (magenta) and either a 4xGCN4-Myo10MD (**Ci**) or 24xGCN4-Myo10MD (**Cii**) linker construct (green).
