## Supplementary figures and images for "Molecular counting of myosin force generators in growing filopodia"

### Fitz et al Video S1

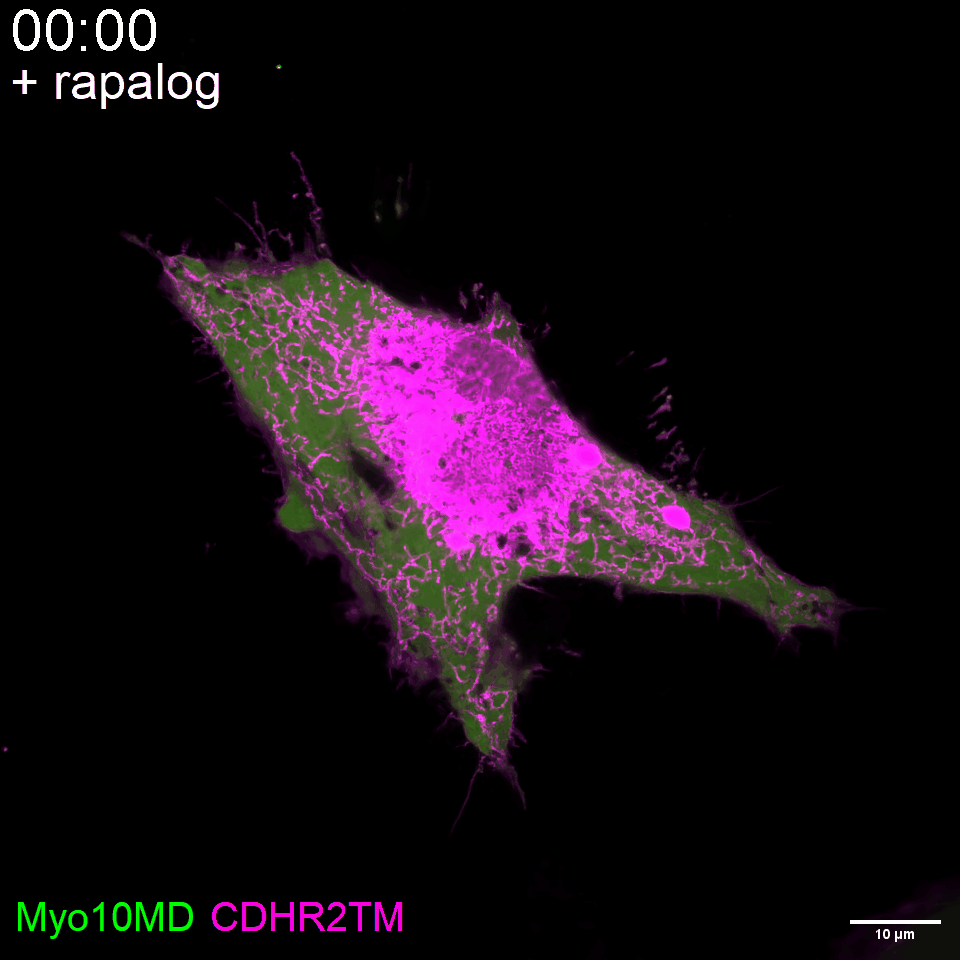

### Fitz et al Video S2

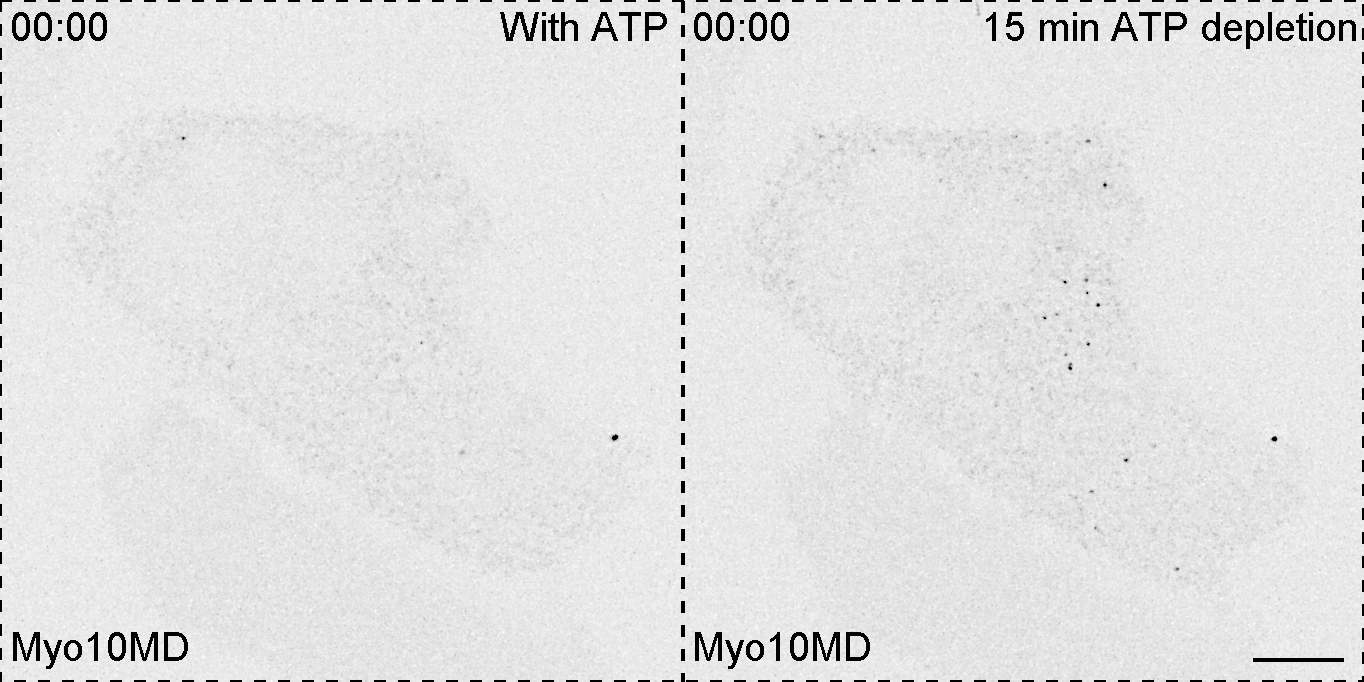
